# REAP: A platform to identify autoantibodies that target the human exoproteome

**DOI:** 10.1101/2021.02.11.430703

**Authors:** Eric Y. Wang, Yile Dai, Connor E. Rosen, Monica M. Schmitt, Mei X. Dong, Elise M. N. Ferré, Feimei Liu, Yi Yang, Jaime A. Gonzalez-Hernandez, Eric Meffre, Monique Hinchcliffe, Fotios Koumpouras, Michail S. Lionakis, Aaron M. Ring

## Abstract

Autoantibodies that recognize extracellular proteins (the “exoproteome”) exert potent biological effects but have proven challenging to detect with existing screening technologies. Here, we developed Rapid Extracellular Antigen Profiling (REAP) as a technique for comprehensive, high-throughput discovery of exoproteome-targeting autoantibodies. With REAP, patient samples are applied to a genetically-barcoded library containing 2,688 human extracellular proteins displayed on the surface of yeast. Antibody-coated cells are isolated by magnetic selection and deep sequencing of their barcodes is used to identify the displayed antigens. To benchmark the performance of REAP, we screened 77 patients with autoimmune polyendocrinopathy candidiasis ectodermal dystrophy (APECED). REAP sensitively and specifically detected known autoantibody reactivities in APECED in addition to numerous previously unidentified reactivities. We further screened 106 patients with systemic lupus erythematosus (SLE) and identified novel autoantibody reactivities against a diverse set of antigens including growth factors, extracellular matrix components, cytokines, and immunomodulatory proteins. Several of these responses were associated with disease severity and specific clinical manifestations of SLE and exerted potent functional effects on cell signaling *ex vivo*. These findings demonstrate the utility of REAP to atlas the expansive landscape of exoproteome-targeting autoantibodies and their impacts on patient health outcomes.

## Introduction

Autoantibodies play a major etiological role across a wide range of diseases spanning autoimmunity, cancer, metabolic dysfunction, cardiovascular disease, infectious diseases, and even neurological and neurodegenerative conditions^1–8^. Though autoantibodies are commonly associated with adverse effects, they can also exhibit disease-ameliorating functions that are beneficial to patients. For example, immunosuppressive anti-cytokine autoantibodies are associated with less severe disease in numerous autoimmune conditions^9, 10^; similarly, anti-tumor specific and opsonizing antibodies are associated with better survival in cancer patients^11–13^. Thus, analogous to genetic mutations, autoantibodies may explain a significant fraction of the clinical and phenotypic variation seen between individuals. Discovery of novel functional autoantibody responses in patients therefore has the potential to uncover key etiologic factors and therapeutic targets similar to the study of human genetics.

Within the human proteome, a particularly important group of autoantibody targets are extracellular and secreted proteins (collectively, the “exoproteome”). Because antibodies are themselves large (150 kDa) secreted proteins, they are most likely to recognize and act upon targets that reside within the same extracellular compartment^14^. While state-of-the-art technologies such as protein/peptide microarrays, proteome-scanning libraries using phage (PhIP-seq) and bacterial display have enabled the discovery of novel autoantibodies in a variety of diseases^15–23^, these systems have limited sensitivity to detect autoantibodies against extracellular targets. This is due in part to the inherent difficulty of working with extracellular proteins, which often have unique folding requirements that include signal peptide removal, disulfide bond formation, and post-translational modifications such as glycosylation. Many of these features are not captured by platforms that express proteins or peptides in prokaryotic systems. Similarly, technologies that rely on the use of peptide fragments are not able to detect autoantibodies that recognize “conformational” protein epitopes (*i.e.*, three dimensional epitopes present when a protein is folded into its native state). This limitation may significantly hamper autoantibody detection, since as many as 90% of antibodies recognize conformational epitopes as opposed to linear peptides^24^.

Here, we describe Rapid Exoproteome Antigen Profiling (REAP), a new method to discover functional antibodies against the exoproteome. REAP leverages yeast-display technology to assess the presence of autoantibody responses to 2,688 extracellular proteins present in patient serum or plasma samples through a next-generation sequencing-based approach. We use REAP to screen a cohort of 77 APECED patients and successfully identify known autoantibodies along with novel “public” (present in many patients) and “private” (present in only a few patients) reactivities. We further apply REAP to a cohort of 106 patients with SLE and identify autoantibodies targeting cytokines, cytokine receptors, growth factors, extracellular matrix components, and immunomodulatory cell surface proteins, and validate several of these reactivities through orthogonal assays. In both SLE and APECED, we identify autoantibody responses that are associated with disease severity and specific clinical disease manifestations. Finally, we find that autoantibodies in SLE patients that target the co-inhibitory ligand PD-L2 and the cytokine IL-33 have functional antagonist activity *ex vivo*. These results indicate that REAP is broadly useful for the discovery of autoantibodies targeting the exoproteome and that functional autoantibodies within patient populations may provide key insights into disease pathogenesis and therapeutic approaches.

## Results

### Development of Rapid Exoproteome Antigen Profiling

To develop a system capable of detecting autoantibody responses against conformational extracellular proteins, we elected to use yeast surface display to comprehensively sample the human exoproteome (**Fig. 1a**). As eukaryotic cells, yeast contain several features that enable them to express extracellular proteins, including endoplasmic reticulum chaperones, glycosylation machinery, and disulfide bond proofreading systems^25^. Accordingly, a diverse range of mammalian extracellular protein families have been successfully expressed with yeast display, including proteins with folds such as the immunoglobulin superfamily (IgSF), TNF superfamily (TNFSF), TNF receptor superfamily (TNFRSF), von Willebrand factor A (vWFA) domains, fibronectin domain, leucine-rich repeat (LRR), EGF-like, insulin-like, cytokines, growth factors, and even complicated assemblies like peptide:MHC complexes, T cell receptors, and intact antibodies^26–41^. We therefore constructed a genetically-barcoded yeast-displayed exoproteome library of approximately 2,700 human extracellular and secreted proteins. The library comprises actively displayed proteins from a wide range of protein families and encompasses 87% of all human exoproteins with extracellular regions from 50-600 amino acids in length (**Fig. 1b, Supplementary Fig. 1a-c**). While there is within-library heterogeneity in individual protein abundance and the number of unique barcodes associated with each gene, the library is relatively uniform and the vast majority of proteins fall within a narrow range suited to coverage by standard next-generation sequencing approaches (**Fig. 1c,d**). Full details on the design and composition of the library are described in the Methods and in **Supplemental Table 1**.

**Figure 1:**
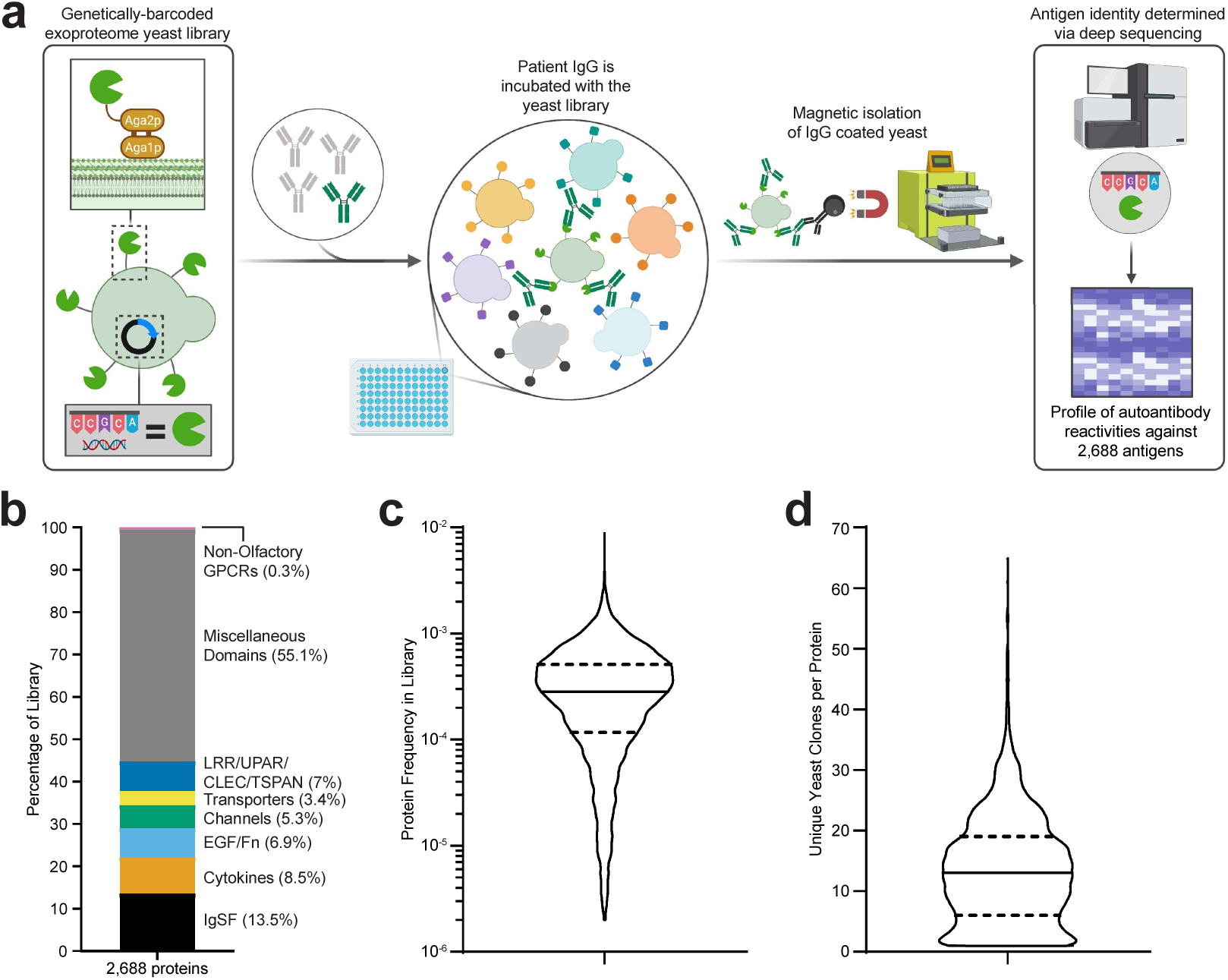
Yeast library and REAP development. **a,** Simplified schematic of REAP. Antibodies are incubated with a genetically-barcoded yeast library displaying members of the exoproteome in 96-well microtiter plates. Antibody bound yeast are enriched by magnetic column-based sorting and enrichment is quantified by next-generation sequencing. **b,** Composition of proteins in the yeast library, categorized by broad protein families. Abbreviations are as follows: immunoglobulin superfamily (IgSF), epidermal growth factor (EGF), fibronectin (Fn), leucine-rich repeat (LRR), urokinase receptor (UPAR), c-type lectin (CLEC), tetraspanin (TSPAN). The cytokine family consists of proteins belonging to tumor necrosis factor, interferon, interleukin, and growth factor protein families. **c & d,** Distribution of total protein frequencies (**c**) and unique yeast clones per protein in the yeast library (**d**). Solid lines indicate the median of the distribution and dotted lines indicate first and third quartiles.

We next optimized procedures for high-throughput identification of seroreactivities against proteins in our exoproteome library for REAP (**Fig. 1d**). Briefly, IgG purified from patient serum or plasma is incubated with the yeast library. Autoantibody-coated cells are then isolated by magnetic separation and deep sequencing of the library-encoded DNA barcodes is used to identify the corresponding antigens encoded by these cells. To quantify the degree of antibody reactivity to a given antigen, we developed a custom scoring algorithm (“REAP Score”) based on the enrichment of each antigen’s barcodes after selection (see Methods). Screening of the exoproteome library with a set of nine conformation-specific monoclonal antibodies against a variety of extracellular proteins showed that all antibody targets were detected specifically and robustly (**Fig. 2a,b**). To further assess the conformational nature of proteins displayed in the library, we performed a REAP screen using a panel of 30 recombinant proteins with known binding partners in the library. REAP accurately detected the cognate binding partners for each of these proteins, with minimal enrichment of off-target proteins (**Fig. 2c**).

**Figure 2:**
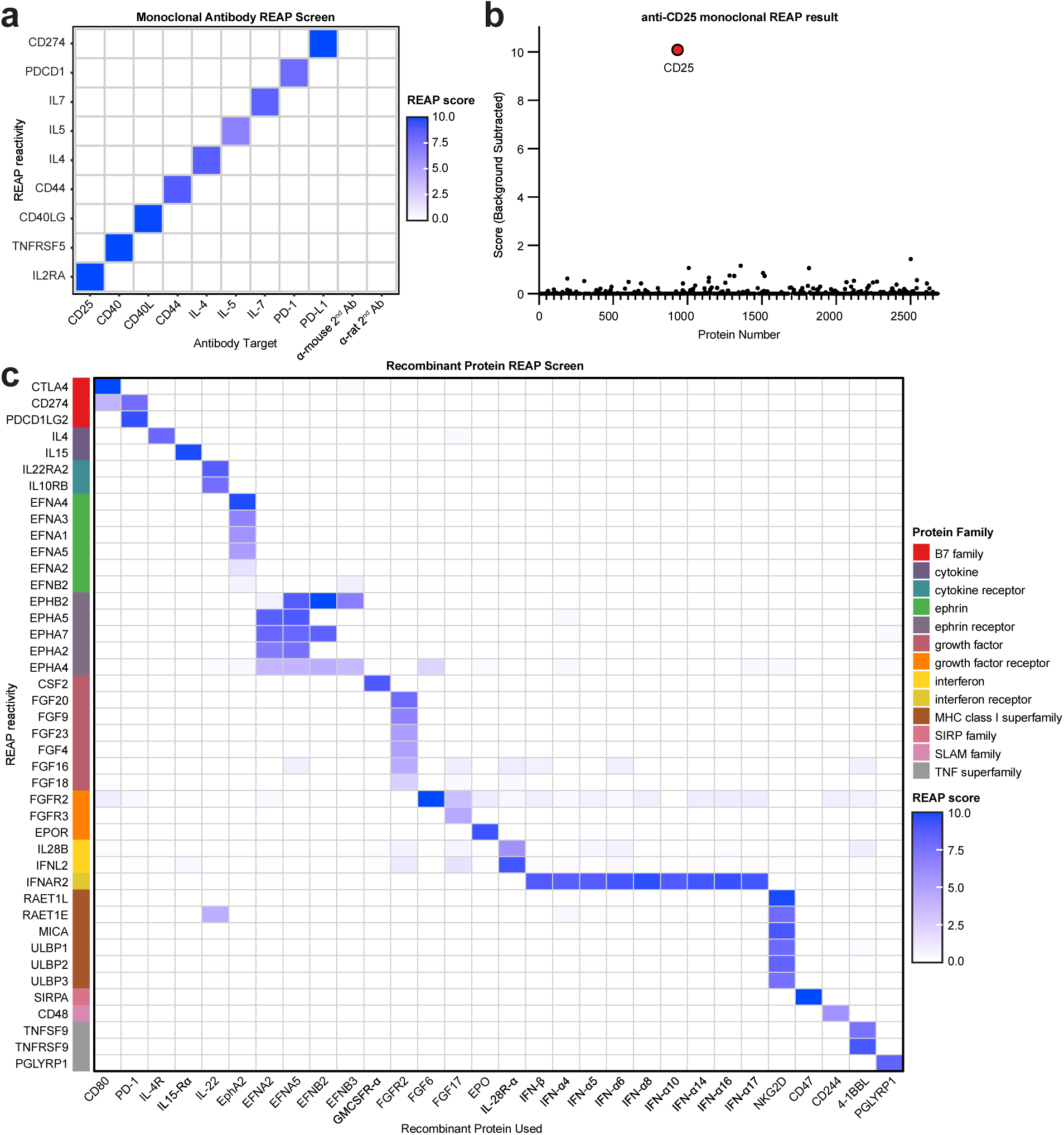
Validation of REAP. A panel of nine monoclonal antibodies were screened using REAP. **a,** Heatmap of results from REAP screen of nine monoclonal antibodies. Only relevant monoclonal antibody targets (gene names) are displayed. **b,** Representative sample from the screen. Monoclonal antibody target is highlighted in red and labelled. Background subtraction was performed by subtracting the score of a selection performed with beads and secondary alone. Scores below the average background level are not shown. **c,** REAP screen performed using recombinant protein in place of IgG.

### Evaluation of REAP performance in APECED

To evaluate the capacity of REAP to detect exoproteome-directed autoantibodies in complex patient samples such as polyclonal responses in serum, we screened a cohort of 77 APECED patients (**Supplementary Table 2**). APECED, also known as autoimmune polyglandular syndrome type-1 (APS-1), is a rare genetic autoimmune disease caused by mutations in the autoimmune regulator (*AIRE*) gene, resulting in loss of central tolerance and the development of chronic mucocutaneous candidiasis (CMC), severe endocrinopathies and other nonendocrine autoimmune sequelae such as pneumonitis, hepatitis, alopecia, vitiligo, and vitamin B12 deficiency/pernicious anemia^42^. Interestingly, APECED patients harbor widespread and pathognomonic autoantibodies targeting numerous cytokines including type I and type III interferons, IL-22, IL-17A, and IL-17F^43–47^. REAP readily identified autoantibody responses against these cytokines in APECED patient samples, but not in samples from healthy controls (**Fig. 3a**). Furthermore, the frequencies of these autoreactivities in APECED patients closely matched the frequencies determined from previous reports using gold-standard methodologies such as enzyme-linked immunosorbent assay (ELISA) and luciferase immunoprecipitation system immunoassay (LIPS) (**Fig. 3b**)^18, 43, 47^. We also identified autoantibodies against gastric intrinsic factor (GIF), lipocalin-1 (LCN1), IL-5, IL-6, protein disulfide-isomerase-like protein of the testis (PDILT), and BPI fold containing family member 1 and 2 (BPIFA1/2), which have been previously described in APECED^18–21, 48^. With respect to GIF reactivities, the results seen with REAP demonstrated strong concordance with anti-GIF ELISA results from the same patients (**Fig. 3c**).

**Figure 3:**
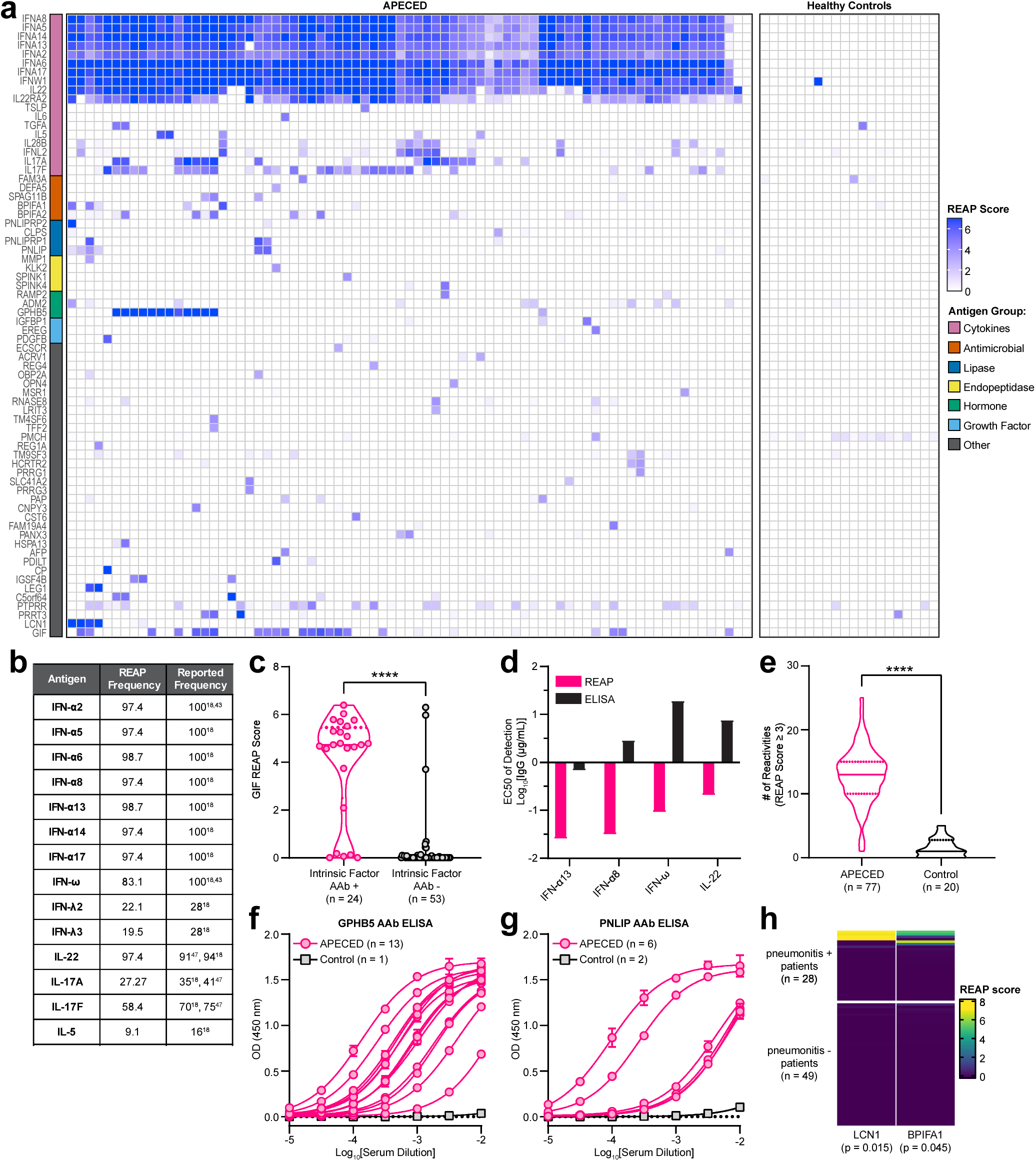
REAP screen of APECED patients. A cohort of 77 APECED patients and 20 healthy controls were screened using REAP. **a,** Heatmap of REAP scores. Antigen groups were manually categorized. **b,** Frequencies of positive reactivities (score ≥ healthy donor average score plus 3 standard deviations) against 14 antigens based on REAP and prior literature^18, 43, 47^. **c,** Violin plot of GIF REAP scores in APECED samples stratified by intrinsic factor clinical autoantibody test results. **d,** EC50 of fitted REAP and ELISA dose response curves for detection of autoantibodies against four proteins in one APECED patient. See **supplementary figure 1e,f** for dose response curves. **e,** Violin plot of the number of reactivities in APECED and control samples at a score cutoff of 3. **f,** anti-GPHB5 and **g,** anti-PNLIP pan-IgG ELISAs conducted with serial dilutions of serum. Error bars represent standard deviation. **h,** Heatmap of LCN1 and BPIFA1 REAP scores in APECED samples stratified by pneumonitis positivity. Listed p-values represent significance for the association between LCN1 or BPIFA1 REAP positivity and pneumonitis. Significance in **c** and **e** was determined using a two-sided Mann-Whitney U test. Significance in **h** was determined using a Fisher Exact Test, where LCN1 and BPIFA1 positivity was defined by a REAP score ≥ 3. In all heatmaps in this figure, score was artificially capped at 7 to aid visualization. In all violin plots in this figure, solid lines represent the median and dotted lines represent the first or third quartile. ****P ≤ 0.0001

To investigate the reproducibility of REAP, we compared log_2_[fold enrichment] between technical (intra-assay) replicates across all APECED patient samples and found strong positive correlations between replicates (median R^2^ = 0.914; **Supplementary Fig. 1d**). To investigate the sensitivity of REAP, we titrated varying amounts of IgG and performed REAP and ELISA side-by-side for four autoantigens (**Supplementary Fig. 1e,f**). In each case, REAP exhibited higher sensitivity than ELISA by 1-2 orders of magnitude, as seen by the calculated EC_50_ values (**Fig. 3d**). Taken in aggregate, these data indicate that REAP is capable of detecting known autoantibody responses against extracellular proteins with high sensitivity and precision.

### APECED patients exhibit broad exoproteome-targeting autoantibody reactivities

Previous reports using protein microarrays and PhIP-seq have shown that APECED patients have greatly elevated numbers of autoantibody reactivities at a proteome-scale compared to healthy controls. Analyzing the REAP data, we found that global autoreactivity present in APECED also extends to the exoproteome (**Fig. 3e, Supplementary Fig. 2a**). While some of the reactivities we observed have been previously characterized, the screen also uncovered numerous previously undescribed “public” (present in more than one patient) and “private” (present in only one patient) reactivities. Two notable public reactivities were those against glycoprotein hormone beta-5 (GPHB5), a thyrostimulin subunit, and pancreatic triacylglycerol lipase (PNLIP), a tissue-restricted antigen that is regulated by AIRE in the thymus^49^. Using ELISA, we confirmed the presence of autoantibody responses against these proteins and found that the titers of autoantibodies were high, ranging from EC50s of approximately 1:100 to 1:10,000 (**Fig. 3f,g**). We additionally were able to correlate particular serological responses to specific, variable clinical features of APECED. For example, we found that autoantibodies against lipocalin-1 (LCN1) and BPIFA1, which had previously been identified in APECED patients with Sjogren’s-like syndrome^48^, were enriched in a subset of APECED patients with pneumonitis (6 out of 28 with pneumonitis), a life-threatening non-endocrine complication of APECED, but universally negative in 49 patients without pneumonitis or healthy controls (**Fig. 3h**). Of note, BPIFA1 reactivity was detected in a patient with biopsy-proven pneumonitis without reactivity to the known lung-targeted autoantibodies KCNRG and BPIFB1, which have an overall sensitivity of ∼75% but are negative in a quarter of patients with biopsy-proven pneumonitis^50^. Interestingly, the single patient in our cohort with exocrine pancreatic insufficiency, a rare manifestation of APECED^42^, uniquely harbored reactivity to colipase (CLPS), an essential cofactor for pancreatic lipase and related lipases (**Fig. 3a**)^51^. Thus, REAP enabled the detection of novel autoantibody reactivities in the monogenic disease APECED, as well as correlations of autoantibodies with clinical features of the disease.

### REAP identifies previously undescribed autoantibody reactivities in SLE patients

We sought to apply REAP to study SLE, a systemic polygenic autoimmune disease characterized by loss of tolerance to nucleic acids^52^. Though autoantibodies are a defining feature in SLE, particularly those against nucleic acids and nuclear protein complexes^53^, the role of functional autoantibodies that target the exoproteome is less well established. We thus performed REAP analysis on samples from a cohort of 106 SLE patients and 20 healthy controls. Patient and control demographics are shown in **Supplementary Table 3**. Compared to APECED, we found that exoproteome-targeting autoantibodies in SLE patients were strikingly heterogeneous; though a wide variety of autoantigens were identified, there were essentially no public autoantigens and most reactivities were present in only a few patients (**Fig. 4a**). Several reactivities identified by REAP included autoantigens that have previously been described in SLE such as IL-6, type I interferons, IL-1α, and TNFα (including identification of a therapeutic anti-TNF antibody administered to one of the patients). We further identified numerous novel autoantibodies targeting other cytokines (*e.g.*, IL-4, IL-33), chemokines (*e.g.*, CXCL3, CCL8), growth factors (*e.g.*, VEGF-B, FGF-21), extracellular matrix components (*e.g.*, epiphycan, vitrin), and immunoregulatory cell surface proteins (*e.g.*, FAS, PD-L2, B7-H4).

**Figure 4:**
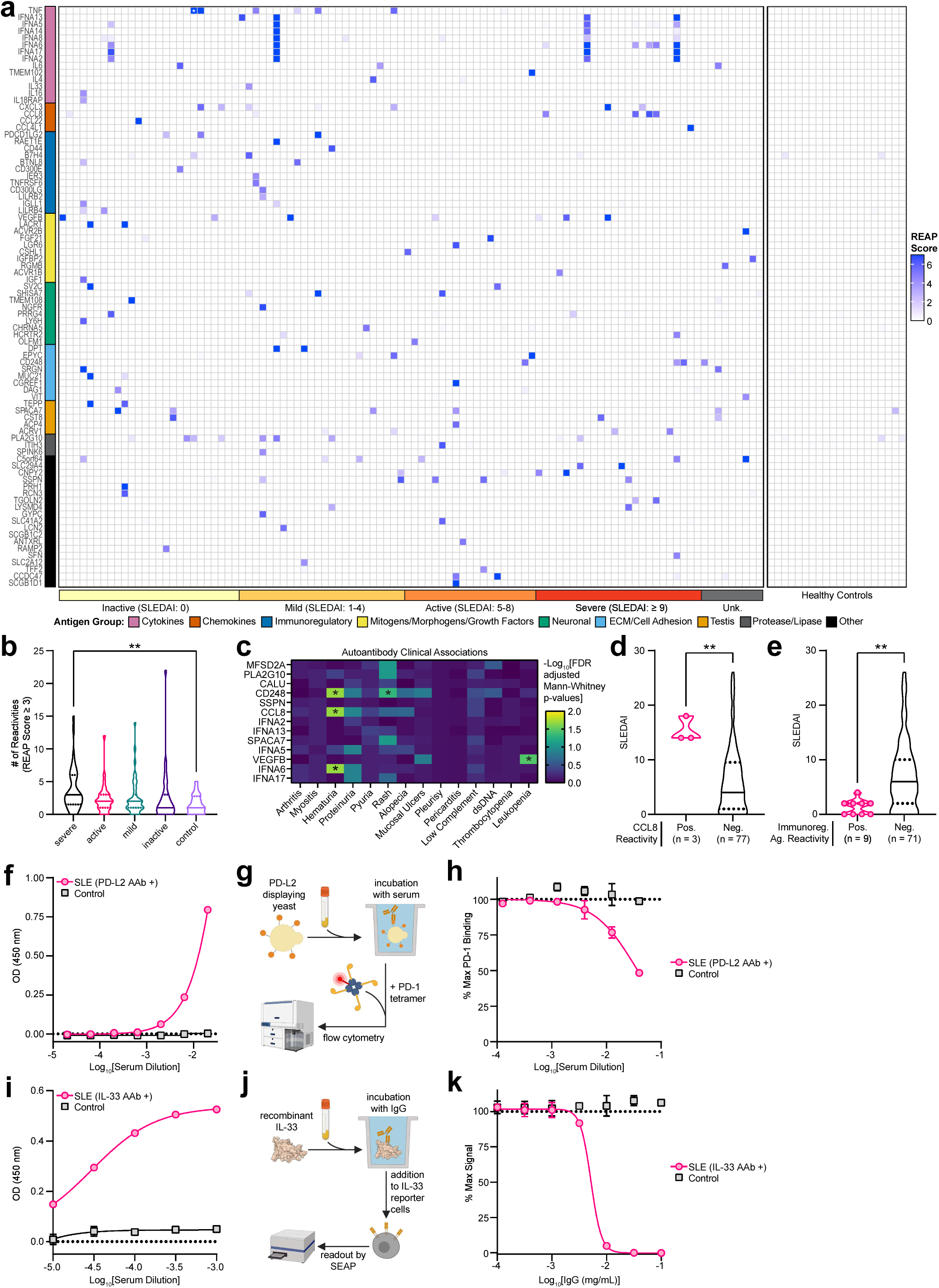
REAP screen of SLE patients. A cohort of 106 unique SLE patients spanning 155 samples and 20 healthy controls was screened using REAP. **a**, Heatmap of REAP scores where each column is a unique patient. For patients with longitudinal samples, the maximum REAP score for each given reactivity is shown. Antigen groups were manually categorized. Patients are ordered from left to right by increasing SLEDAI score. White stars symbolize detection of a therapeutic antibody. Score was artificially capped at 7 to aid visualization. **b,** Violin plots of the number of reactivities in SLE samples stratified by disease severity and control samples at a score cutoff of 3. Significance was determined using a Kruskal-Wallis test followed by a Dunnett’s test. **c,** Heatmap of false discovery rate-adjusted p-values from two-sided Mann-Whitney U tests comparing REAP score distributions for specific proteins between patients stratified by disease manifestations. Only reactivities positive in at least 3 patients were tested. **d,** SLEDAI scores for SLE patients stratified by reactivity against CCL8. **e,** SLEDAI scores for SLE patients positive or negative by REAP score for reactivities against immunoregulatory antigens (defined in **a**). **f,** anti-PD-L2 and **i,** anti-IL-33 pan-IgG ELISAs conducted with serial dilutions of SLE or control serum. **g,** schematic and **h,** results of PD-L2 blocking assay conducted with serial dilutions of serum from a control and the SLE patient in **f**. **j,** schematic and **k,** results of IL-33 neutralization assay conducted with serial dilutions of IgG from a control and the SLE patient in **i**. Significance in **d** and **e** was determined using a two-sided Mann-Whitney U test. All error bars in this figure represent standard deviation. For all analyses in this figure, positive reactivities were defined as those with REAP score ≥ 3. *P ≤ 0.05, **P ≤ 0.01.

To validate the large number of candidate autoantibody reactivities identified by REAP, we tested autoantibody reactivities against several different proteins using LIPS and ELISA and subsequently confirmed 16 of these autoantigens (**Table 1,** **Fig. 4f,i, Supplementary Fig. 3a-h, j, n-r**). The subset of confirmed autoantibody reactivities consisted of both shared and private reactivities and included examples of potentially pathological and well as immunomodulatory reactivities, such as those against the extracellular matrix component epiphycan (**Supplementary Fig. 3n**), the cytokine receptor IL-18Rβ **(Supplementary Fig. 3p**), the death receptor FAS/TNFRSF6 (**Supplementary Fig. 3e**), the co-inhibitory ligand PD-L2 (**Fig. 4f**), and the IL-1 family cytokine IL-33 (**Fig. 4i**). We additionally characterized the titers and IgG isotypes for several of these responses, finding that they spanned a wide range of titers (1:10 to >1:10,000) and isotype classes (**Supplementary Fig. 3n-r, t-v**). Using these results, as well as orthogonal validations of known APECED reactivities (**Supplementary Fig. 3i-m**), we performed receiver operating characteristic (ROC) analysis to quantify the performance of the REAP scoring algorithm. We found that REAP score sensitively and specifically predicted autoantibody reactivity by ELISA and/or LIPS, with an area under the curve (AUC) of 0.892 **(Supplementary Fig. 3s**). Because REAP exhibits greater sensitivity for some antigens than the ELISA/LIPS “gold standards” (as was the case for type I IFN autoantibodies in APECED), this number may represent a conservative estimate of the true performance of REAP in predicting autoantibody reactivity.

**Table 1:**
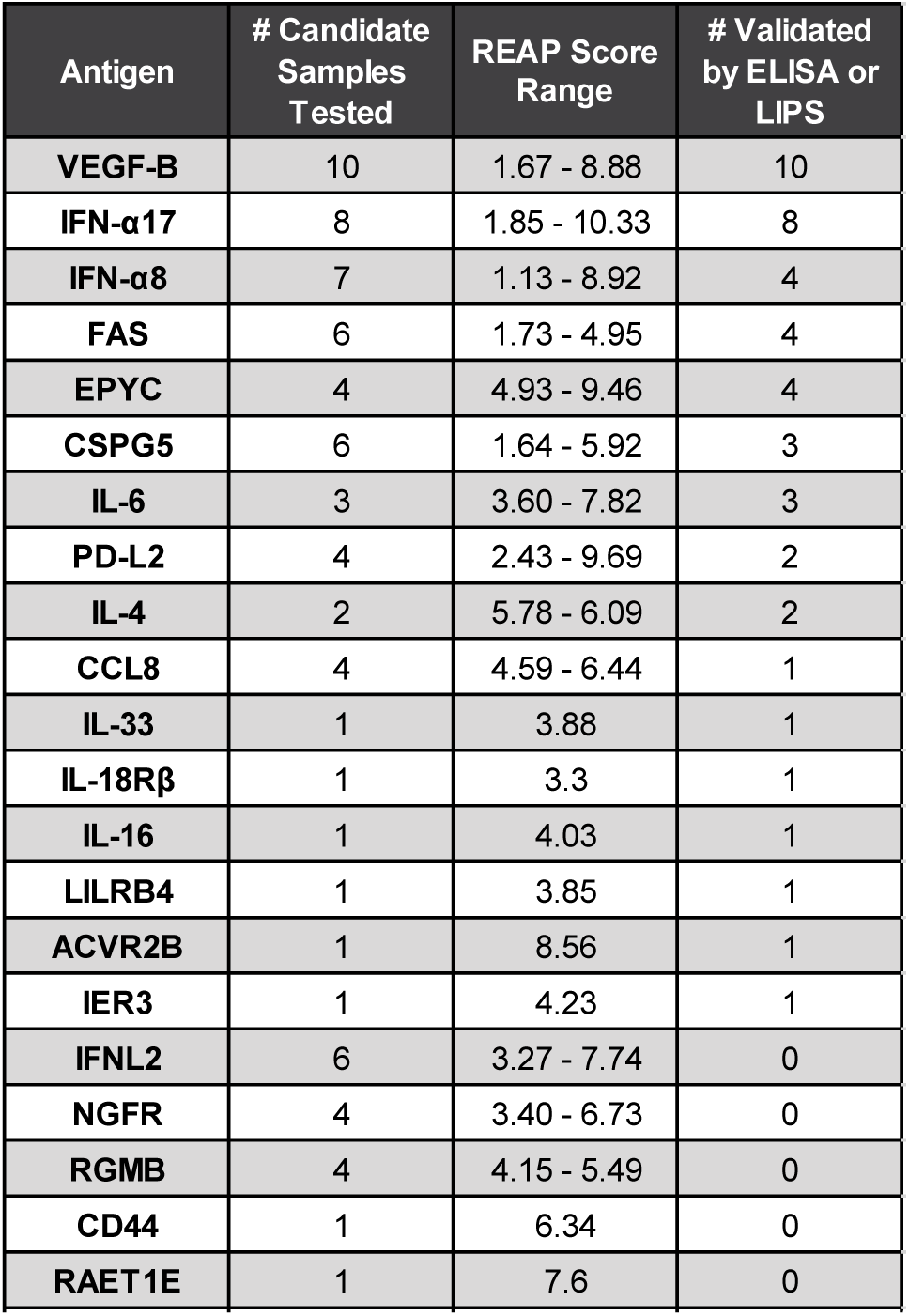
Orthogonal validation of SLE autoantibody reactivities identified in REAP.

### Exoproteome-targeting autoantibodies in SLE are functional and correlate with disease severity

Given the broad distribution of autoantibody responses in SLE, we wondered if particular responses or patterns of reactivity were associated with specific clinical features of the disease. At a global level, we found that the total numbers of autoantibody reactivities identified with REAP correlated with worse clinical severity, as measured by the Systemic Lupus Erythematosus Disease Activity Index (SLEDAI) score^54^. In particular, we found that samples from patients with severe disease (SLEDAI score ≥ 9) had significantly increased numbers of autoantibodies compared to healthy controls (**Fig. 4b, Supplementary Fig. 2b**). Furthermore, SLE patients in all severity groups had reactivities that were not observed in healthy individuals and these patterns of reactivity were associated with particular SLE disease phenotypes. For instance, we found that autoantibody reactivities against the chemokine CCL8, the cytokine IFN-alpha-6, and the C-type lectin CD248 (endosialin) were significantly associated with hematuria and that VEGF-B reactivities were associated with leukopenia (**Fig. 4c**). Additionally, patients positive for CCL8 reactivity had significantly higher SLEDAI scores, indicating more severe disease (**Fig. 4d**). By contrast, patients who exhibited autoreactivity against a set of immunoregulatory proteins (PD-L2, RAET1E, CD44, B7H4, BTNL8, CD300E, IER3, TNFRSF6, CD300LG, LILRB2, IGLL1, and LILRB4) had significantly lower SLEDAI scores compared to patients negative for these autoantibodies (**Fig. 4e**).

Finally, we characterized the functionality of autoantibodies against two novel autoantigens identified by REAP, PD-L2 and IL-33. As the primary biological function of PD-L2 is mediated by its binding to its receptor PD-1, we tested whether autoantibodies against PD-L2 could block this interaction. Serum samples from an SLE patient with anti-PD-L2 autoantibodies were present at titers >1:100 and inhibited the interaction between PD-L2 and PD-1 in a dose-dependent manner, while serum from a control patient without anti-PD-L2 autoantibodies did not (**Fig. 4f-h**). To test the functional effects of anti-IL-33 autoantibodies, we used a HEK-Blue IL-33 reporter cell line, which produces secreted alkaline phosphatase downstream of an NFκB promoter that is activated by the IL-33 pathway. Bulk IgG (isolated via protein G) from the SLE patient harboring anti-IL-33 autoantibodies potently neutralized IL-33 signaling with an IC_50_ less than 0.01 mg/mL, while IgG from a control patient without anti-IL-33 autoantibodies had no neutralizing effect (**Fig. 4i-k**). These findings underscore the ability of REAP to discover novel autoantibodies with functional biological effects.

## Discussion

In the present study we show that REAP is a sensitive and high-throughput platform for discovery of exoproteome-directed autoantibodies. By querying antigens in a conformationally-active state, REAP enables identification of autoantibodies that are difficult to detect, if not entirely invisible to other technologies. This was particularly evident in our screen of APECED samples, as we found that REAP was considerably more accurate in detecting a well-defined subset of known extracellular autoantigen reactivities compared to protein arrays and phage-peptide display approaches. Furthermore, REAP enabled the identification of numerous previously undescribed autoantigens in APECED patients, a surprising finding given how extensively autoantibodies have been studied in this patient population.

We also identified a large set of previously undescribed autoantibody reactivities against the exoproteome in SLE patients, a considerably more heterogeneous population than APECED. The vast majority of these novel autoreactivities were relatively private with a prevalence of <5% and in some cases present in only a single patient. Though these autoantibody responses are rare, our studies suggest that they can exert large biological effects that could meaningfully impact disease progression, akin to the effect of rare genetic variants. For example, we identified a single SLE patient with mild disease activity (SLEDAI score of 1) who had extraordinarily high-titer autoantibodies against IL-33 that potently neutralized IL-33 signaling *in vitro*. This suggests that these IL-33 antibodies may have played a protective role that ameliorated the severity of the disease in this individual and, by extension, that IL-33 blockade could represent a potential therapeutic strategy in SLE. Indeed, circulating IL-33 concentrations are elevated in SLE patients and are positively correlated with C-reactive protein concentrations and clinical manifestations such as thrombocytopenia and erythrocytopenia^55, 56^. Similarly, preclinical studies in mouse models have demonstrated that IL-33 exposure is associated with autoantibody production and that neutralization of IL-33 suppresses lupus-like disease^57, 58^. Beyond IL-33, we also found that SLE patients with autoreactivity against a set of immunoreceptors had substantially lower disease severity, indicating that disruption of those pathways and/or opsonization of cells that express the receptors could similarly exert a protective effect. Future investigation is warranted to determine the prevalence of these autoantibodies in SLE patients and their potential protective effects on a larger, confirmatory cohort. Nevertheless, our finding that functional autoantibodies responses are highly variable between patients underscores the need for technologies like REAP that can provide comprehensive, unbiased antibody profiling for large numbers of patients. Without sufficient sample throughput and representation of the exoproteome, these rare, but impactful autoantibody responses might not be readily detected.

REAP does have important limitations. While our data indicate that most exoproteome antigens are displayed on the surface of yeast and we additionally demonstrated that dozens of the library members are biochemically active (via recapitulating known binding interactions), not all members of the exoproteome can be expressed in the yeast system. This may be due to lack of specific chaperones, expression partners, or post-translational modifications required for protein folding and activity. Furthermore, while yeast do perform O- and N- linked glycosylation, their glycosylation patterns are characterized by a hypermannose structure that is highly divergent from glycosylation seen in humans^59^. Thus, autoantibodies recognizing specific glycoforms of their antigens would not be detected with REAP. Further improvement in the REAP platform could therefore involve yeast strain engineering to co-express mammalian chaperone proteins to enhance folding of human antigens and glycosylation enzymes to produce more human-like glycosylation patterns, as has been described for the yeast species *Pichia pastoris*^60^.

Though we initially applied REAP to the study of autoimmune conditions, an intriguing avenue of future study with REAP and other serological profiling technologies is to characterize autoantibody responses in diseases such as cancer, infectious diseases, and neurological conditions that are not considered to have a primarily autoimmune etiology. Identification of disease-modifying antibody responses in such conditions could implicate new molecular pathways that contribute to disease pathology as well as novel therapeutic targets and molecular diagnostics. Furthermore, patient autoantibodies could represent potential therapeutic agents themselves. Technologies such as REAP can enable these discoveries by revealing the diverse landscape of functional autoantibody responses that influence health and disease.

## Materials and Methods

### Library production

#### Library design

An initial library of 3093 human extracellular proteins was assembled based on protein domains, immunological functions, and yeast-display compatibility. The extracellular portion of each protein was identified by manual inspection of topological domains annotated in the SwissProt database (January 2018). For proteins with uncertain topology, full sequences were run through SignalP 4, Topcons, and GPIPred to identify most likely topologies. For proteins with multiple extracellular portions, in general the longest individual region was chosen for initial amplification. cDNAs for chosen proteins were purchased from GE Dharmacon or DNASU. The protein sequences were further modified to match isoforms available in purchased cDNAs. An inventory of antigens included in the library are compiled in supplementary table 1.

#### Library construction

A two-step PCR process was used to amplify cDNAs for cloning into a barcoded yeast-display vector. cDNAs were amplified with gene-specific primers, with the forward primer containing a 5’ sequence (CTGTTATTGCTAGCGTTTTAGCA) and the reverse primer containing a 5’ sequence (GCCACCAGAAGCGGCCGC) for template addition in the second step of PCR. PCR reactions were conducted using 1 µL pooled cDNA, gene-specific primers, and the following PCR settings: 98 ⁰C denaturation, 58 ⁰C annealing, 72 ⁰C extension, 35 rounds of amplification. 1 µL of PCR product was used for direct amplification by common primers Aga2FOR and 159REV, and the following PCR settings: 98 ⁰C denaturation, 58 ⁰C annealing, 72 ⁰C extension, 35 rounds of amplification. PCR product was purified using magnetic PCR purification beads (AvanBio). 90 µL beads were added to the PCR product and supernatant was removed. Beads were washed twice with 200 µL 70% ethanol and resuspended in 50 µL water to elute PCR products from the beads. Beads were removed from purified PCR products. The 15bp barcode fragment was constructed by overlap PCR. 4 primers (bc1, bc2, bc3, bc4; sequences listed below) were mixed in equimolar ratios and used as a template for a PCR reaction using the following PCR settings: 98 ⁰C denaturation, 55 ⁰C annealing, 72 ⁰C extension, 35 rounds of amplification. Purified product was reamplified with the first and fourth primer using identical PCR conditions. PCR products were run on 2% agarose gels and purified by gel extraction (Qiagen). Purified barcode and gene products were combined with linearized yeast-display vector (pDD003 digested with EcoRI and BamHI) and electroporated into JAR300 yeast using a 96-well electroporator (BTX Harvard Apparatus) using the following electroporation conditions: Square wave, 500 V, 5 ms pulse, 2 mm gap. Yeast were immediately recovered into 1 mL liquid synthetic dextrose medium lacking uracil (SDO - Ura) in 96-well deep well blocks and grown overnight at 30°C. Yeast were passaged once by 1:10 dilution in SDO-Ura, then frozen as glycerol stocks. To construct the final library, 2.5 µL of all wells were pooled and counted. A limited dilution of 300,000 clones was sub-sampled and expanded in SDO-Ura. Expression was induced by passaging into synthetic galactose medium lacking uracil (SGO-Ura) at a 1:10 dilution and growing at 30°C overnight. 10^8^ yeast were pelleted and resuspended in 1 mL PBE (PBS with 0.5% BSA and 0.5 mM EDTA) containing 1:100 anti-FLAG PE antibody (BioLegend). Yeast were stained at 4° for 75 minutes, then washed twice with 1 mL PBE and sorted for FLAG display on a Sony SH800Z cell sorter. Sorted cells were expanded in SDO-Ura supplemented with 35 µg/mL chloramphenicol, expanded, and frozen as the final library.

**bc1**- TTGTTAATATACCTCTATACTTTAACGTCAAGGAGAAAAAACCCCGGATC
**bc2**- CTGCATCCTTTAGTGAGGGTTGAANNNNNNNNNNNNNNNTTCGATCCGGGGTTTTTTCTCCTTG
**bc3**- TTCAACCCTCACTAAAGGATGCAGTTACTTCGCTGTTTTTCAATATTTTCTGTTATTG C
**bc4**- TGCTAAAACGCTAGCAATAACAGAAAATATTGAAAAACAGCG

#### Barcode identification

Barcode-gene pairings were identified using a custom Tn5-based sequence approach. Tn5 transposase was purified as previously described, using the on-column assembly method for loading oligos^61^. DNA was extracted from the yeast library using Zymoprep-96 Yeast Plasmid Miniprep kits or Zymoprep Yeast Plasmid Miniprep II kits (Zymo Research) according to standard manufacturer protocols. 5 µL of purified plasmid DNA was digested with Tn5 in a 20 µL total reaction as previously described. 2 µL of digested DNA was amplified using primers index1 and index2, using the following PCR settings: 98 ⁰C denaturation, 56 ⁰C annealing, 72 ⁰C extension, 25 rounds of amplification. The product was run on a 2% gel and purified by gel extraction (Qiagen). Purified product was amplified using primers index3 and index4, using the following PCR settings: 98 ⁰C denaturation, 60 ⁰C annealing, 72 ⁰C extension, 25 rounds of amplification. In parallel, the barcode region alone was amplified using primers index1 and index5, using the following PCR settings: 98 ⁰C denaturation, 56 ⁰C annealing, 72 ⁰C extension, 25 rounds of amplification. The product was run on a 2% gel and purified by gel extraction (Qiagen). Purified product was amplified using primers index3 and index6, using the following PCR settings: 98 ⁰C denaturation, 60 ⁰C annealing, 72 ⁰C extension, 20 rounds of amplification. Both barcode and digested fragment products were run on a 2% gel and purified by gel extraction (Qiagen). NGS library was sequenced using an Illumina MiSeq and Illumina v3 MiSeq Reagent Kits with 150 base pair single-end sequencing according to standard manufacturer protocols. Gene-barcode pairings were identified using custom code. Briefly, from each read, the barcode sequence was extracted based on the identification of the flanking constant vector backbone sequences, and the first 25 bp of sequence immediately following the constant vector backbone-derived signal peptide were extracted and mapped to a gene identity based on the first 25 bp of all amplified cDNA constructs. The number of times each barcode was paired with an identified gene was calculated. Barcode-gene pairings that were identified more than twice, with an overall observed barcode frequency of greater than .0002% were compiled. For barcodes with multiple gene pairings matching the above criteria, the best-fit gene was manually identified by inspection of all barcode-gene pairing frequencies and, in general, identification of the most abundant gene pairing. In the final library, 2,688 genes were confidently mapped to 35,835 barcodes.

### Rapid Extracellular Antigen Profiling

#### Antibody purification and yeast adsorption

20 µL protein G magnetic resin (Lytic Solutions) was washed twice with 100 µL sterile PBS, resuspended in 50 µL PBS, and added to 50 µL serum or plasma. Serum-resin mixture was incubated for three hours at 4 ⁰C with shaking. Resin was washed five times with 200 µL PBS, resuspended in 90 µL 100 mM glycine pH 2.7, and incubated for five minutes at room temperature. Supernatant was extracted and added to 10 µL sterile 1M Tris pH 8.0 (purified IgG). Empty vector (pDD003) yeast were expanded in SDO-Ura at 30 ⁰C. One day later, yeast were induced by 1:10 dilution in SGO-Ura for 24 hours. 10^8^ induced yeast were washed twice with 200 µL PBE (PBS with 0.5% BSA and 0.5 mM EDTA), resuspended with 100 µL purified IgG, and incubated for three hours at 4 ⁰C with shaking. Yeast-IgG mixtures were placed into 96 well 0.45 um filter plates (Thomas Scientific) and yeast-depleted IgG was eluted into sterile 96 well plates by centrifugation at 3000 g for 3 minutes.

#### Antibody yeast library selections

Transformed yeast were expanded in SDO-Ura at 30 ⁰C. One day later, at an optical density (OD) below 8, yeast were induced by resuspension at an OD of 1 in SGO-Ura supplemented with ten percent SDO-Ura and culturing at 30⁰C for 20 hours. Prior to selection, 400 µL pre-selection library was set aside to allow for comparison to post-selection libraries. 10^8^ induced yeast were washed twice with 200 µL PBE and added to wells of a sterile 96-well v-bottom microtiter plate. Yeast were resuspended in 100 µL PBE containing appropriate antibody concentration and incubated with shaking for 1 hour at 4 ⁰C. Unless otherwise indicated, 10 μg antibody per well was used for human serum or plasma derived antibodies and 1 μg antibody was used for monoclonal antibodies. Yeast were washed twice with 200 µL PBE, resuspended in 100 µL PBE with a 1:100 dilution of biotin anti-human IgG Fc antibody (clone HP6017, BioLegend) for human serum or plasma derived antibodies or a 1:25 dilution of biotin goat anti-rat or anti-mouse IgG antibody (A16088, Thermo Fisher Scientific; A18869, Thermo Fisher Scientific) for monoclonal antibodies. Yeast-antibody mixtures were incubated with shaking for 30 minutes at 4 ⁰C. Yeast were washed twice with 200 µL PBE, resuspended in 100 µL PBE with a 1:20 dilution of Streptavidin MicroBeads (Miltenyi Biotec), and incubated with shaking for 30 minutes at 4 ⁰C. Yeast were then pelleted and kept on ice. Multi-96 Columns (Miltenyi Biotec) were placed into a MultiMACS M96 Separator (Miltenyi Biotec) and the separator was placed into positive selection mode. All following steps were carried out at room temperature. Columns were equilibrated with 400 µL 70% ethanol followed by 700 µL degassed PBE. Yeast were resuspended in 200 µL degassed PBE and placed into the columns. After the mixture had completely passed through, columns were washed three times with 700 µL degassed PBE. To elute the selected yeast, columns were removed from the separator and placed over 96-well deep well plates. 700 µL degassed PBE was added to each well of the column and the column and deep well plate were spun at 50 g for 30 seconds. This process was repeated 3 times. Selected yeast were pelleted, and recovered in 1 mL SDO -Ura at 30 ⁰C.

#### Recombinant protein yeast library selections

All pre-selection and yeast induction steps were performed identically as those of the antibody yeast library selections. 10^8^ induced yeast were washed twice with 200 µL PBE and added to wells of a sterile 96-well v-bottom microtiter plate. Yeast were resuspended in 100 µL PBE containing 75 μL clarified protein expression supernatant and incubated with shaking for 1 hour at 4 ⁰C. Yeast were washed twice with 200 µL PBE, resuspended in 100 µL PBE with 5 μL μMACS Protein G MicroBeads (Miltenyi Biotec), and incubated with shaking for 30 minutes at 4 ⁰C. Selection of yeast using the MultiMACS M96 Separator and subsequent steps were performed identically as those of the antibody yeast library selections.

#### Next generation sequencing library preparation and sequencing

DNA was extracted from yeast libraries using Zymoprep-96 Yeast Plasmid Miniprep kits or Zymoprep Yeast Plasmid Miniprep II kits (Zymo Research) according to standard manufacturer protocols. A first round of PCR was used to amplify a DNA sequence containing the protein display barcode on the yeast plasmid. PCR reactions were conducted using 1 µL plasmid DNA, 159_DIF2 and 159_DIR2 primers (sequences listed below), and the following PCR settings: 98 ⁰C denaturation, 58 ⁰C annealing, 72 ⁰C extension, 25 rounds of amplification. PCR product was purified using magnetic PCR purification beads (AvanBio). 45 µL beads were added to the PCR product and supernatant was removed. Beads were washed twice with 100 µL 70% ethanol and resuspended in 25 µL water to elute PCR products from the beads. Beads were removed from purified PCR products. A second round of PCR was conducted using 1 µL purified PCR product, Nextera i5 and i7 dual-index library primers (Illumina), and the following PCR settings: 98 ⁰C denaturation, 58 ⁰C annealing, 72 ⁰C extension, 25 rounds of amplification. PCR products were pooled and run on a 1% agarose gel. The band corresponding to 257 base pairs was cut out and DNA (NGS library) was extracted using a QIAquick Gel Extraction Kit (Qiagen) according to standard manufacturer protocols. NGS library was sequenced using an Illumina MiSeq and Illumina v3 MiSeq Reagent Kits with 75 base pair single-end sequencing or using an Illumina NovaSeq 6000 and Illumina NovaSeq S4 200 cycle kit with 101 base pair paired-end sequencing according to standard manufacturer protocols. A minimum of 50,000 reads per sample was collected and the pre-selection library was sampled at ten times greater depth than other samples.

**159_DIF2**- TCGTCGGCAGCGTCAGATGTGTATAAGAGACAGNNNNNNNNNNGAGAAAAAACCCCGGATCG
**159_DIR2**- GTCTCGTGGGCTCGGAGATGTGTATAAGAGACAGNNNNNNNNNNACGCTAGCAATAACAGAAAATATTG

#### Data analysis

REAP scores were calculated as follows. First, barcode counts were extracted from raw NGS data using custom codes and counts from technical replicates were summed. Next, aggregate and clonal enrichment was calculated using edgeR^62^ and custom codes. For aggregate enrichment, barcode counts across all unique barcodes associated with a given protein were summed, library sizes across samples were normalized using default edgeR parameters, common and tagwise dispersion were estimated using default edgeR parameters, and exact tests comparing each sample to the pre-selection library were performed using default edgeR parameters. Aggregate enrichment is thus the log2 fold change values from these exact tests with zeroes in the place of negative fold changes. Log2 fold change values for clonal enrichment were calculated in an identical manner, but barcode counts across all unique barcodes associated with a given protein were not summed. Clonal enrichment for a given reactivity was defined as the fraction of clones out of total clones that were enriched (log2 fold change ≥ 2). Aggregate *E_a_*) and clonal enrichment *E_c_*) for a given protein, a scaling factor *β_u_*) based on the number of unique yeast clones (yeast that have a unique DNA barcode) displaying a given protein, and a scaling factor *β_f_*) based on the overall frequency of yeast in the library displaying a given protein were used as inputs to calculate the REAP score, which is defined as follows.

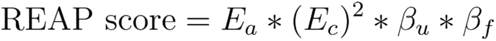

*β_u_* and *β_f_* are logarithmic scaling factors that progressively penalize the REAP score of proteins with low numbers of unique barcodes or low frequencies in the library. *β_u_* is applied to proteins with ≤ 5 unique yeast clones in the library and *β_f_* is applied to proteins with a frequency ≤ 0.0001 in the library. *β_f_* was implemented to mitigate spurious enrichment signals from low frequency proteins, which could occur due to sequencing errors or stochasticity in the selection process. *β_u_* was implemented because the clonal enrichment metric is less valid for proteins with low numbers of unique yeast clones, decreasing confidence in the validity of the reactivity. *β_u_* and *β_f_* are defined as follows where is the number of unique yeast clones for a given protein and is the log10 transformed frequency of a given protein in the library.

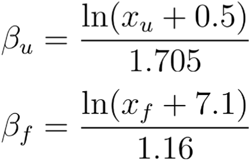

### Recombinant protein production

#### REAP recombinant protein production

Proteins were produced as human IgG1 Fc fusions to enable binding of secondary antibody and magnetic beads to the produced proteins during the REAP process. Sequences encoding the extracellular portions of proteins-of-interests that were present in the yeast display library were cloned by Gibson assembly into a modified pD2610-v12 plasmid (ATUM). Modifications include addition of an H7 signal sequence followed by a (GGGGS)_3_ linker and a truncated human IgG1 Fc (N297A). Protein-of-interest sequences were inserted directly downstream of the H7 leader sequence. Protein was produced by transfection into Expi293 cells (Thermo Fisher Scientific) in 96-well plate format. One day prior to transfection, cells were seeded at a density of 2 million cells per mL in Expi293 Expression Medium (Thermo Fisher Scientific). In a 96-well plate, 0.5 μg plasmid DNA was diluted added to 25 μL Opti-MEM (Thermo Fisher Scientific) and mixed gently. In a separate 96-well plate, 1.35 μL ExpiFectamine was added to 25 μL Opti-MEM and mixed gently. The ExpiFectamine-Opti-MEM mixture was added to the diluted DNA, mixed gently, and incubated for 20 minutes at room temperature. Expi293 cells were diluted to a density of 2.8 million cells per mL and 500 μL of cells were added to each well of a 96-well deep well plate. 50 μL of the DNA-ExpiFectamine-Opti-MEM mixture was added to each well. The plate was sealed with Breathe-Easier sealing film (Diversified Biotech) and incubated in a humidified tissue culture incubator (37 ⁰C, 8% CO_2_) with shaking at 1,200 rpm so that cells were kept in suspension. 18-20 hours post-transfection, 25 μL enhancer 2 and 2.5 μL enhancer 1 (Thermo Fisher Scientific) were added to each well. 4 days post-transfection, media was clarified by centrifugation at 3000-4000 g for 5 minutes. Clarified media was used for recombinant protein REAP.

#### ELISA protein production

Sequences encoding the extracellular portions of proteins-of-interests that were present in the yeast display library were cloned by Gibson assembly into pEZT_Dlux, a modified pEZT-BM vector. The pEZT-BM vector was a gift from Ryan Hibbs (Addgene plasmid #74099). Modifications included insertion of an H7 Leader Sequence followed by an AviTag (Avidity), HRV 3C site, protein C epitope, and an 8x his tag. Protein-of-interest sequences were inserted directly downstream of the H7 leader sequence. Protein was produced by transfection into Expi293 cells (Thermo Fisher Scientific) according to standard manufacturer protocols. Transfected cells were maintained according to manufacturer protocols. 4 days post-transfection, media was clarified by centrifugation at 300 g for 5 minutes. Protein was purified from clarified media by nickel-nitrilotriacetic acid (Ni-NTA) chromatography and desalted into HEPES buffered saline + 100 mM sodium chloride, pH 7.5. Protein purity was verified by SDS-PAGE.

#### Biotinylated protein production

Sequences encoding the extracellular portions of proteins-of-interests were cloned into pEZT_Dlux as described above. Protein was expressed and purified as described above minus desalting. Enzymatic biotinylation with BirA ligase was performed and protein was purified by size-exclusion fast protein liquid chromatography using a NGC Quest 10 Chromatography System (Bio-Rad).

#### LIPS protein production

Sequences encoding Lucia luciferase (InvivoGen) fused by a GGSG linker to the N-terminus of the protein-of-interest extracellular portion (as defined above) were cloned by Gibson assembly into pEZT-BM. Protein was produced by transfection into Expi293 cells (Thermo Fisher Scientific) according to standard manufacturer protocols. Transfected cells were maintained according to manufacturer protocols. 3 days post-transfection, media was clarified by centrifugation at 300 g for 5 minutes. Clarified media was used in luciferase immunoprecipitation systems assays.

### Enzyme-linked immunosorbent assays (ELISAs)

200 or 400 ng of purchased or independently produced recombinant protein in 100 µL of PBS pH 7.0 was added to 96-well flat bottom Immulon 2HB plates (Thermo Fisher Scientific) and placed at 4 ⁰C overnight. Plates were washed once with 225 µL ELISA wash buffer (PBS + 0.05% Tween 20) and 150 µL ELISA blocking buffer (PBS + 2% Human Serum Albumin) was added to the well. Plates were incubated with shaking for 2 hours at room temperature. ELISA blocking buffer was removed from the wells and appropriate dilutions of sample serum in 100 µL ELISA blocking buffer were added to each well. Plates were incubated with shaking for 2 hours at room temperature. Plates were washed 6 times with 225 µL ELISA wash buffer and 1:5000 goat anti-human IgG HRP (Millipore Sigma) or anti-human IgG isotype specific HRP (Southern Biotech; IgG1: clone HP6001, IgG2: clone 31-7-4, IgG3: clone HP6050, IgG4: clone HP6025) in 100 µL ELISA blocking buffer was added to the wells. Plates were incubated with shaking for 1 hour at room temperature. Plates were washed 6 times with 225 µL ELISA wash buffer. 50 µL TMB substrate (BD Biosciences) was added to the wells and plates were incubated for 15 minutes (pan-IgG ELISAs) or 20 minutes (isotype specific IgG ELISAs) in the dark at room temperature. 50 µL 1 M sulfuric acid was added to the wells and absorbance at 450 nm was measured in a Synergy HTX Multi-Mode Microplate Reader (BioTek).

### Luciferase immunoprecipitation systems (LIPS) assays

Pierce Protein A/G Ultralink Resin (5 µL; Thermo Fisher Scientific) and 1 µL sample serum in 100 µL Buffer A (50 mM Tris, 150 mM NaCl, 0.1% Triton X-100, pH 7.5) was added to 96-well opaque Multiscreen HTS 96 HV 0.45 um filter plates (Millipore Sigma). Plates were incubated with shaking at 300 rpm for 1 hour at room temperature. Supernatant in wells was removed by centrifugation at 2000 g for 1 minute. Luciferase fusion protein (10^6^ RLU) was added to the wells in 100 µL Buffer A. Plates were incubated with shaking at 300 rpm for 1 hour at room temperature. Using a vacuum manifold, wells were washed 8 times with 100 µL Buffer A followed by 2 washes with 100 µL PBS. Remaining supernatant in wells was removed by centrifugation at 2000 g for 1 minute. Plates were dark adapted for 5 minutes. An autoinjector equipped Synergy HTX Multi-Mode Microplate Reader (BioTek) was primed with QUANTI-Luc Gold (InvivoGen). Plates were read using the following per well steps: 50 µL QUANTI-Luc Gold injection, 4 second delay with shaking, read luminescence with an integration time of 0.1 seconds and a read height of 1 mm.

### PD-L2 blocking assay

A single clone of PD-L2 displaying yeast was isolated from the library and expanded in SDO-Ura at 30 ⁰C. Yeast were induced by 1:10 dilution into SGO-Ura and culturing at 30 ⁰C for 24 hours. 10^5^ induced PD-L1 yeast were washed twice with 200 μL PBE and added to wells of a 96-well v-bottom microtiter plate. Yeast were resuspended in 25 μL PBE containing serial dilutions of sample serum and incubated with shaking for 1 hour at 4 ⁰C. PD-1 tetramers were prepared by incubating a 5:1 ratio of biotinylated PD-1 and PE streptavidin (BioLegend) for 10 minutes on ice in the dark. Yeast were washed twice with 200 μL PBE, resuspended in 25 μL PBE containing 10 nM previously prepared PD-1 tetramers, and incubated with shaking for 1 hour at 4 ⁰C. Yeast were washed twice with 200 μL PBE and resuspended in 75 μL PBE. PE fluorescent intensity was quantified by flow cytometry using a Sony SA3800 Spectral Cell Analyzer. Percent max binding was calculated based on fluorescent PD-1 tetramer binding in the absence of any serum.

### IL-33 neutralization assay

#### IL-33 reporter cell line construction

The full-length coding sequence for ST2 was cloned by Gibson assembly into the lentiviral transfer plasmid pL-SFFV.Reporter.RFP657.PAC, a kind gift from Benjamin Ebert (Addgene plasmid #61395). HEK-293FT cells were seeded into a 6-well plate in 2 mL growth media (DMEM with 10% (v/v) FBS, 100 units/mL penicillin, and 0.1 mg/mL streptomycin) and were incubated at 37°C, 5% CO2. Once cells achieved 70-80% confluence approximately one day later, cells were transfected using TransIT-LT1 (Mirus Bio) in Opti-MEM media (Life Technologies). TransIT-LT1 Reagent was pre-warmed to room temperature and vortexed gently. For each well, 0.88 ug lentiviral transfer plasmid along with 0.66 ug pSPAX2 (Addgene plasmid #12260) and 0.44 ug pMD2.G (Addgene plasmid #12259), kind gifts from Didier Trono, were added to 250 μL Opti-MEM media and mixed gently. TransIT-LT1 reagent (6 μl) was added to the DNA mixture, mixed gently, and incubated at room temperature for 15-20 minutes. The mixture was added dropwise to different areas of the well. Plates were incubated at 37°C, 5% CO2; 48hrs later, the virus-containing media was collected and filtered with a 0.45μm low protein-binding filter. HEK-Blue IL-18 cells (InvivoGen) were seeded into a 6-well plate in 1 mL growth media (DMEM with 10% (v/v) FBS, 100 units/mL penicillin, and 0.1 mg/mL streptomycin) and 1 mL virus-containing media. Cells were incubated at 37°C, 5% CO2 for two days before the media was changed.

#### Reporter cell stimulation and reading

Purified IgG titrations and 2 nM IL-33 were mixed in 50 µL assay media (DMEM with 10% (v/v) FBS, 100 units/mL penicillin, and 0.1 mg/mL streptomycin) and incubated with shaking for 1 hour at room temperature. Approximately 50,000 IL-33 reporter cells in 50 µL assay media were added to wells of a sterile tissue culture grade flat-bottom 96-well plate. IgG-IL-33 mixtures were added to respective wells (1 nM IL-33 final concentration). Plates were incubated at 37°C, 5% CO2 for 20 hours, then 20 µL media from each well was added to 180 μL room temperature QUANTI-Blue Solution (InvivoGen) in a separate flat-bottom 96-well plate and incubated at 37°C for 3 hours. Absorbance at 655 nm was measured in a Synergy HTX Multi-Mode Microplate Reader (BioTek). Percent max signal was calculated based on signal generated by IL-33 in the absence of any serum.

### ROC analysis of REAP score performance

Orthogonal validation data for the receiver operator curve (ROC) analysis was obtained by ELISA, LIPS, or clinical autoantibody tests. For ELISA and LIPS, valid reactivities were defined as those 3 standard deviations above the healthy donor average for a given protein in each assay. ROC analysis was performed using 247 test pairs across 25 different proteins. A full list of ROC inputs can be found in **Supplementary Data 1**.

### Patient Samples

#### SLE patients

Collection of SLE patient blood samples was approved by the Yale Human Research Protection Program Institutional Review Boards (protocol ID 1602017276). All patients met the 2012 SLICC classification criteria for SLE^63^. Clinical information was gathered via retrospective EMR review. Informed consent was obtained from all patients.

#### APECED patients

Collection of APECED patient blood samples was performed under a NIAID IRB-approved prospective natural history study (11-I-0187, NCT01386437). Patients underwent a comprehensive clinical evaluation at the NIH Clinical Center including a detailed history and physical examination, laboratory and radiologic evaluations and consultations by a multidisciplinary team of specialists including infectious disease, immunology, genetics, endocrinology, gastroenterology, hepatology, pulmonology, dermatology, dental, and ophthalmology, as previously described^64^. All study participants provided written informed consent.

### Statistical analysis

Statistical details of experiments can be found in the figure legends. All REAP screens and experimental assays were performed with technical replicates. Data analysis was performed using R, Python, Excel, and GraphPad Prism. Unless otherwise specified, adjustment for false discovery rate was performed using the Benjamini-Hochberg procedure.

## Data Availability

Data are available from the corresponding author upon reasonable request.

## Code Availability

All code will be available at GitHub.

## Figure Legends

**Supplementary Figure 1:**
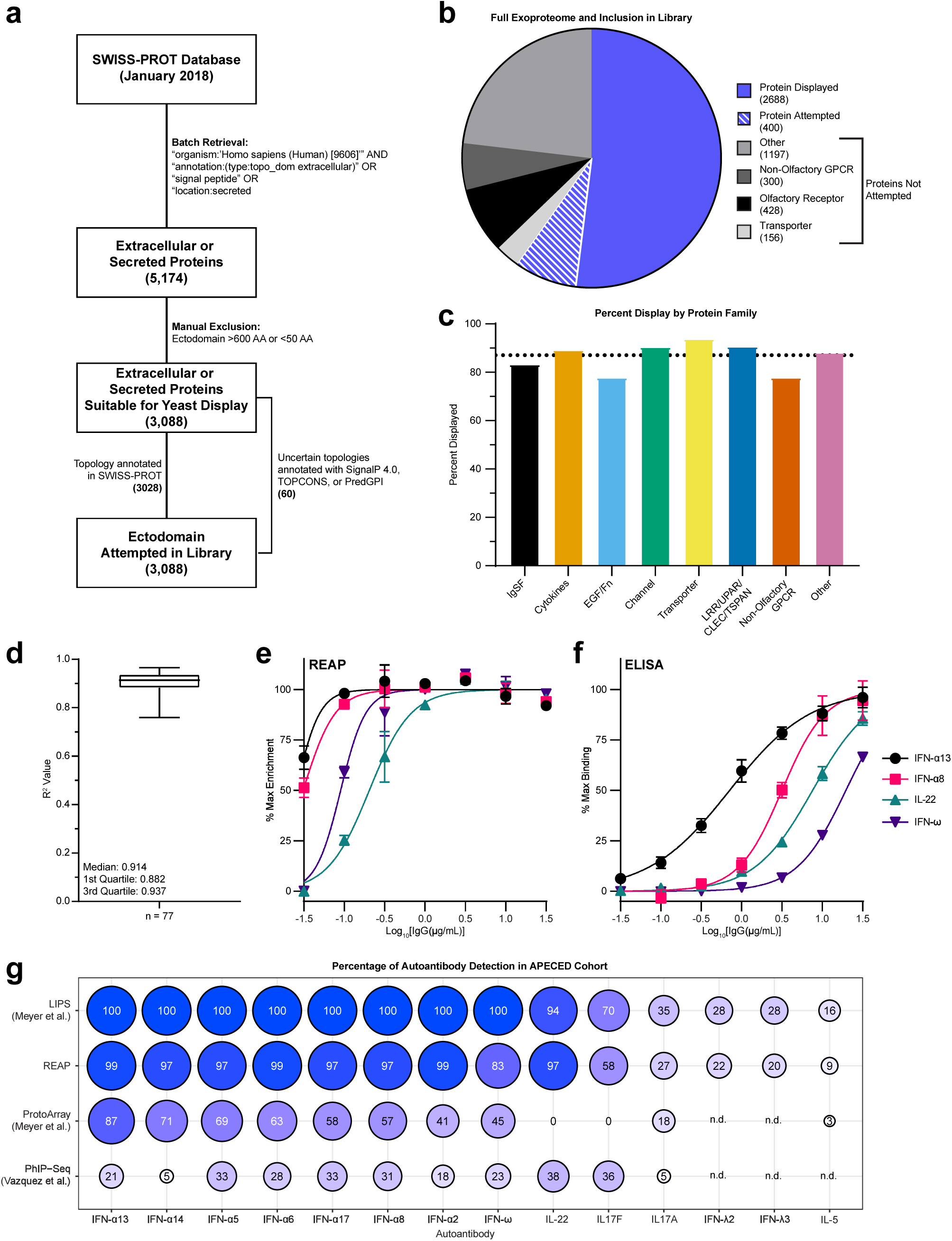
Exoproteome yeast display library properties. **a,** Flowchart of steps in identification and annotation of extracellular or secreted proteins for inclusion in the library. **b,** Pie chart of all extracellular or secreted proteins identified in **a**. Proteins were not attempted if they had an ectodomain less than 50 amino acids or less than 600 amino acids. **c,** Percent of proteins displayed in each protein family included in the library. The dotted line represents the aggregate display level in the library. Abbreviations are as follows: immunoglobulin superfamily (IgSF), epidermal growth factor (EGF), fibronectin (Fn), leucine-rich repeat (LRR), urokinase receptor (UPAR), c-type lectin (CLEC), tetraspanin (TSPAN). The cytokine family consists of proteins belonging to tumor necrosis factor, interferon, interleukin, and growth factor protein families. **d,** Box plot of Log_2_[fold enrichment] *R^2^* coefficient of determination values between technical replicates of APECED patients screened in **figure 2**. **e & f,** REAP (**e**) versus ELISA (**f**) dose-response curve comparison for APECED autoantibodies against four proteins. REAP data is from a screen conducted using varying concentrations of AIRE.19 IgG. Curves were fit using a sigmoidal 4 parameter logistic curve. For REAP, curves were fit based on Log_2_[fold enrichment]. For ELISA, curves were fit based on optical density at 450 nm. Error bars represent standard error of the mean. **g,** Comparison of autoantibody detection frequencies in APECED patient cohorts by REAP, LIPS^18^, ProtoArray^18^, and PhIP-Seq^21^. Frequencies are listed as a percentage inside each circle. Size and color of circles are proportional to detection frequency. For REAP, detection frequency was calculated as in **figure 2b**. For LIPS and ProtoArray, detection frequencies were provided in the corresponding publication. For PhIP-Seq, detection frequency was calculated based on figures in the corresponding publication. For reactivities labelled n.d., either data was not publicly available or the autoantibody was not tested for in the corresponding assay.

**Supplementary Figure 2:**
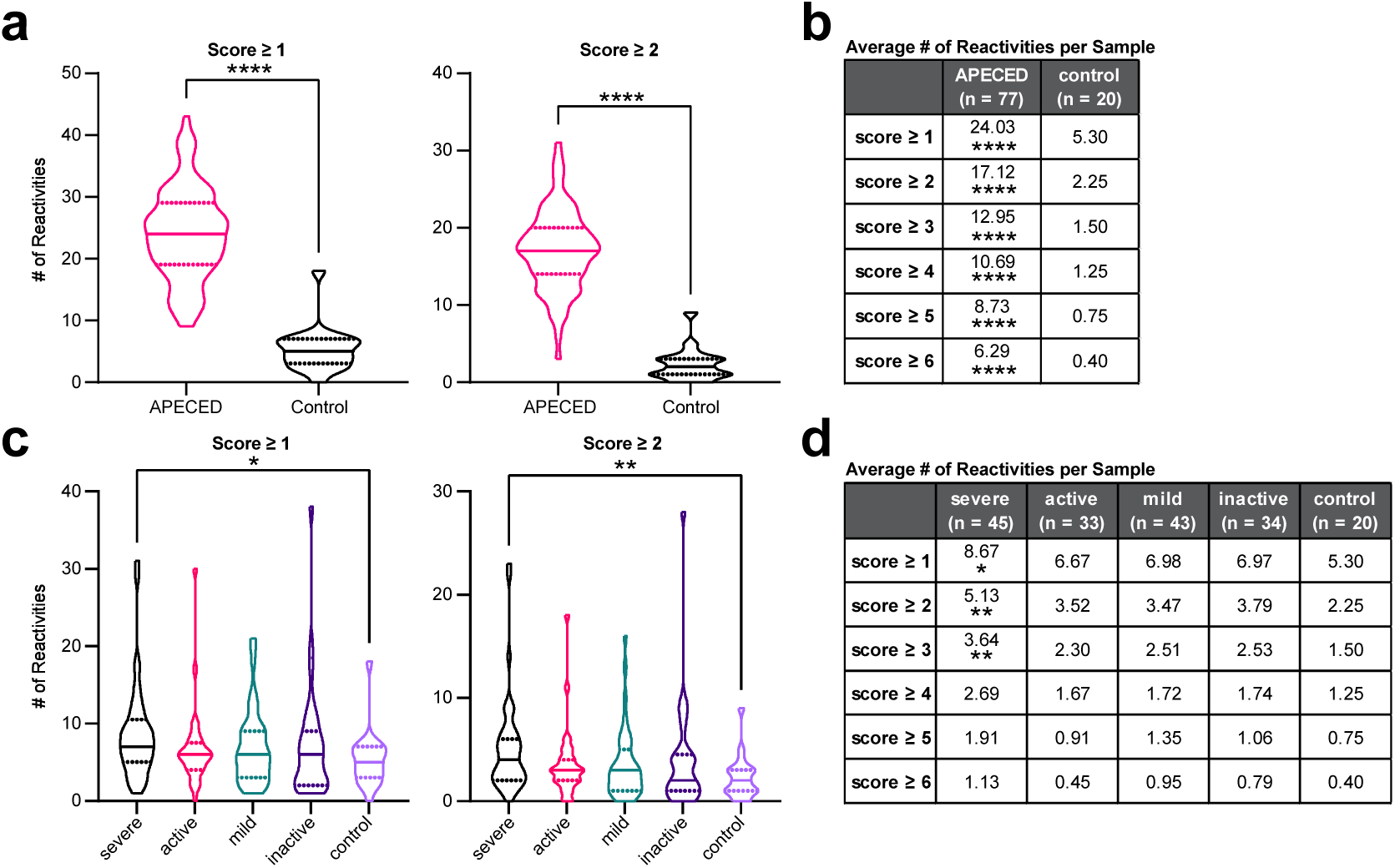
APECED and SLE reactivity distributions. **a,** Violin plots of the number of reactivities in APECED and control samples at a score cutoff of 1 or 2. **b,** Mean number of reactivities in APECED and control samples at various score cutoffs, along with indicators of significance. **c,** Violin plots of the number of reactivities in SLE samples stratified by disease severity and control samples at a score cutoff of 1 or 2. **d,** Mean number of reactivities in SLE samples stratified by disease severity and control samples at various score cutoffs. Comparisons were made between each disease severity group and the control group. Significance in **a** and **b** was calculated using a two-sided Mann-Whitney U test. Significance in **c** and **d** was determined using a Kruskal-Wallis test followed by a Dunnett’s test.

**Supplementary Figure 3:**
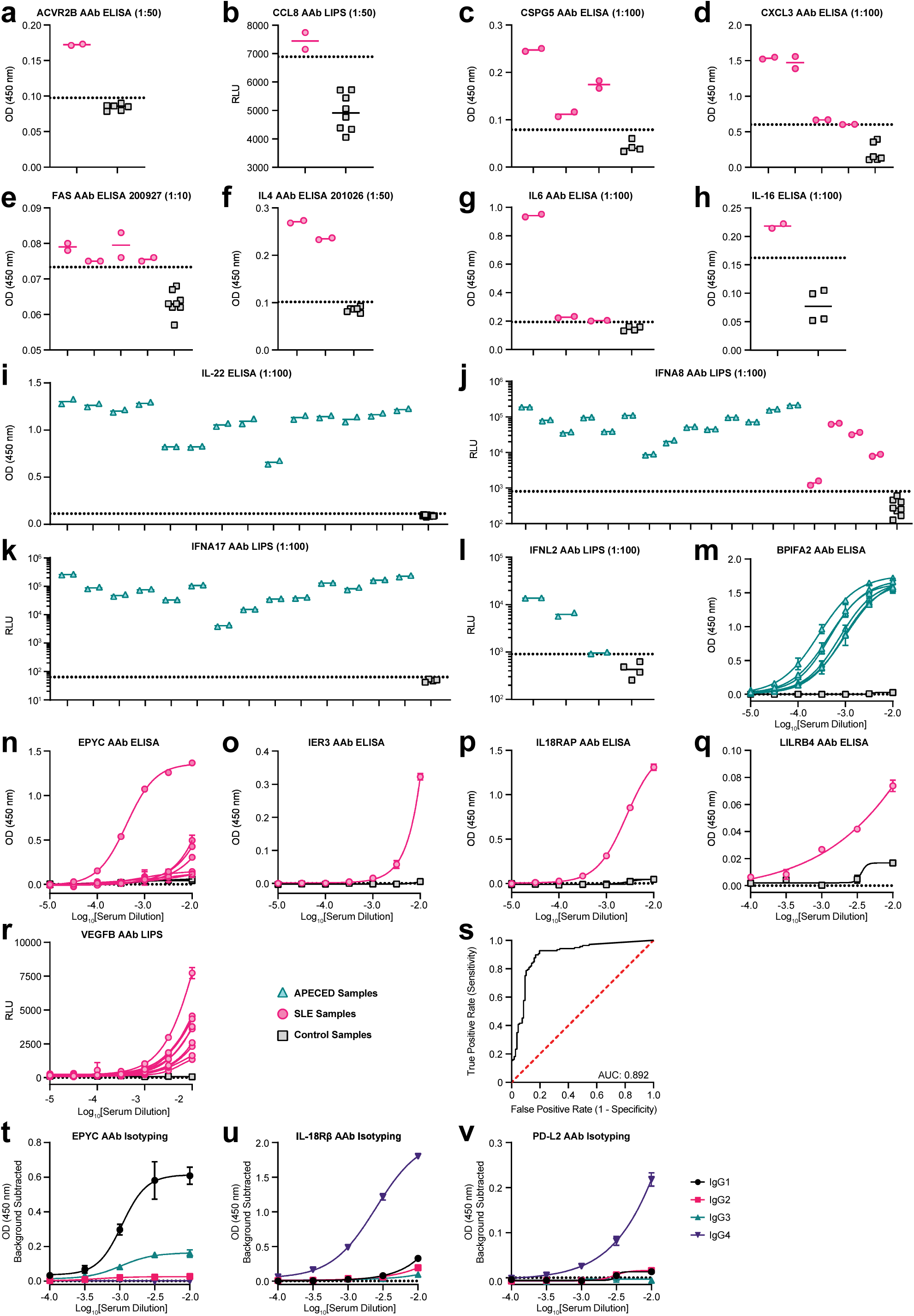
REAP validation and ROC analysis. a-l, Single-point ELISAs or LIPS conducted with SLE, APECED, or control serum to detect autoantibodies against ACVR2B (**a**), CCL8 (**b**), CSPG5 (**c**), CXCL3 (**d**), Fas (**e**), IL-4 (**f**), IL-6 (**g**), IL-16 (**h**), IL-22 (**i**), IFN-α8 (**j**), IFN-α7 (**k**), and IFNL2 (**l**). Serum dilutions are listed in the title of each plot. **m-r,** ELISAs or LIPS conducted with serial dilutions of SLE, APECED, or control serum to detect autoantibodies against BPIFA2 (**m**), EPYC (**n**), IER3 (**o**), IL18RAP (**p**), LILRB4 (**q**), and VEGF-B (**r**). Dotted lines in **a-l** represent the control average + 3 standard deviations. **s,** Receiver operating characteristic curve of the ability of REAP score to predict validation of a REAP reactivity in an orthogonal assay. A full description of this analysis can be found in the materials and methods section. **t,** Anti-epiphycan IgG subclass specific ELISA conducted with serial dilutions of serum from the SLE patient with highest titers in **n**. **u,** Anti-IL-18RAcP subclass specific ELISA conducted with serial dilutions of serum from the SLE patient in **p**. **v,** Anti-PD-L2 IgG subclass specific ELISAs conducted with serial dilutions of serum from the SLE patient in **figure 3f**. All error bars in this figure all represent standard deviation. All curves in this figure were fit using a sigmoidal 4 parameter logistic curve.

Supplementary Table 1: List of protein antigens included in library

**Supplementary Table 2:** APECED patient demographics and clinical characteristics.

| APECED cohort characteristics (n = 77) | Number (%) |
| --- | --- |
| **Age\*** | 24 (14.4) |
| **Gender (female)** | 45 (58) |
| **Ethnicity** |  |
| White Non-Hispanic | 68 (88) |
| White/Hispanic | 5 (7) |
| **AIRE alleles\*\*** |  |
| c.967_979del13 | 79 (51) |
| c.769C>T | 21 (14) |
| **Clinical manifestations** |  |
| Chronic mucocutaneous candidiasis | 66 (86) |
| Adrenal insufficiency | 62 (81) |
| Hypoparathyroidism | 63 (82) |
| Hypothyroidism | 18 (23) |
| Hypogonadism | 26 (34) |
| Autoimmune pneumonitis | 28 (36) |
| Autoimmune hepatitis | 25 (33) |
| Intestinal dysfunction | 53 (69) |
| Exocrine pancreatic insufficiency | 1 (1) |
| Asplenia | 10 (13) |
| Alopecia | 26 (34) |
| Vitiligo | 19 (25) |
| Sjogren's-like syndrome | 30 (39) |
| Autoimmune gastritis | 30 (39) |
| B12 deficiency | 20 (26) |
| Intrinsic factor antibody | 24 (31) |
| **Lung-targeted autoantibodies\*\*\*** |  |
| BPIFB1 | 19 (26) |
| KCNRG | 4 (6) |
| *Age is represented as mean (standard deviation) in years<br>**The denominator for *AIRE* mutant alleles is 154<br>***Data available for 72 patients<br>AIRE, autoimmune regulator; APECED, autoimmune polyendocrinopathy-candidiasis-ectodermal dystrophy; BPIFB1, BPI fold containing family B member 1 |  |

**Supplementary Table 3:** SLE patient and control demographics and clinical characteristics.

| Mean (SD) or as indicated | SLE Cohort<br>(n = 85*) | Healthy Controls<br>(n = 20) |
| --- | --- | --- |
| **Age, (years)** | 41.7 (12.6) | 37.2 (11) |
| **Gender, N (% female)** | 76 (89.4) | 12 (60) |
| **Ethnicity, N (%)** |  |  |
| Hispanic | 22 (26) | 3 (15) |
| Non-Hispanic | 35 (41) | 8 (40) |
| African American | 28 (33) | 9 (45) |
| **Clinical Manifestations, N (%)** |  |  |
| Skin | 40 (47.1) |  |
| Mucocutaneous | 16 (18.8) |  |
| Musculoskeletal | 29 (34.1) |  |
| Renal | 20 (23.5) |  |
| Cardiorespiratory | 4 (4.7) |  |
| Hematological | 7 (8.2) |  |
| Neuropsychiatric | 0 (0) |  |
| **Serologies, N (%)** |  |  |
| Positive dsDNA | 40 (47.1) |  |
| Low complement | 34 (40) |  |
| **SLEDAI score** | 6.3 (6.1) |  |
| **Medications, any use N (%)** |  |  |
| Prednisone | 40 (47.1) |  |
| Hydroxychloroquine | 72 (84.7) |  |
| Mycophenolate mofetil | 24 (28.2) |  |
| Methotrexate | 6 (7.1) |  |
| Azathioprine | 4 (4.7) |  |
| Belimumab | 6 (7.1) |  |
| Others (cyclophosphamide, rituximab, tacrolimus, infliximab, etc.) | 10.6% |  |
| Abbreviations: SLEDAI (Systemic Lupus Erythematosus Disease Activity Index).<br>Prednisone dosing ranges from 5 mg daily to 60 mg daily.<br>*Complete clinical data was not available for a subset of patients. A total of 106 patients were screened. |  |  |

Supplementary Data 1: Receiver operating characteristic analysis inputs

## Acknowledgements

The authors gratefully acknowledge all members of the Ring Laboratory and Daniela Deny for helpful advice and technical assistance. The authors also thank the Yale Section of Rheumatology, Allergy & Immunology for providing systemic lupus erythematosus (SLE) samples from its repository as well as the members of the Yale Rheumatology Clinical and Translational Research Laboratory, Shannon Teaw BS and Michelle Cheng BA for aliquoting SLE patient sera. This work was supported by gifts from the Mathers Family Foundation, the Colton Foundation, the Ludwig Family Foundation, and a supplement to the Yale Cancer Center Support Grant 3P30CA016359-40S4. A.M.R. is additionally supported by an NIH Director’s Early Independence Award (DP5OD023088), a Pew-Stewart Award, and the Robert T. McCluskey Foundation. This work was supported by the Division of Intramural Research of the National Institute of Allergy and Infectious Diseases, NIH.

## Author Contributions

E.Y.W., Y.D., C.E.R., F.L., and Y.Y. performed experiments. M.X.D. provided lupus patient samples and clinical annotations. M.M.S., E.M.N.F., and M.S.L. provided APECED patient samples and clinical annotations. E.Y.W., Y.D., E.M., M.S.L., and A.M.R. analyzed data, M.H., M.S.L., and A.M.R. provided project supervision. E.Y.W. and A.M.R. wrote the paper.

## Competing Interests

E.Y.W., Y.D., C.E.R., and A.M.R. are inventors of a patent describing the REAP technology and A.M.R. is the founder of Seranova Bio.

## Notes

### Summary of Updates

corrected mislabelled y-axis in Figure 3e

## References

1. Ludwig, R. J. et al. Mechanisms of Autoantibody-Induced Pathology. Front. Immunol. 8, 603 (2017).

2. Kazarian, M. & Laird-Offringa, I. A. Small-cell lung cancer-associated autoantibodies: potential applications to cancer diagnosis, early detection, and therapy. Mol. Cancer 10, 33 (2011).

3. Leslie, R. D., Palmer, J., Schloot, N. C. & Lernmark, A. Diabetes at the crossroads: relevance of disease classification to pathophysiology and treatment. Diabetologia 59, 13–20 (2016).

4. Menconi, F., Marcocci, C. & Marinò, M. Diagnosis and classification of Graves’ disease. Autoimmun. Rev. 13, 398–402 (2014).

5. Meier, L. A. & Binstadt, B. A. The Contribution of Autoantibodies to Inflammatory Cardiovascular Pathology. Front. Immunol. 9, 911 (2018).

6. Ercolini, A. M. & Miller, S. D. The role of infections in autoimmune disease. Clin. Exp. Immunol. 155, 1–15 (2009).

7. De Virgilio, A. et al. Parkinson’s disease: Autoimmunity and neuroinflammation. Autoimmun. Rev. 15, 1005–1011 (2016).

8. Britschgi, M. et al. Neuroprotective natural antibodies to assemblies of amyloidogenic peptides decrease with normal aging and advancing Alzheimer’s disease. Proc. Natl. Acad. Sci. U. S. A. 106, 12145–12150 (2009).

9. Cappellano, G. et al. Anti-cytokine autoantibodies in autoimmune diseases. Am. J. Clin. Exp. Immunol. 1, 136–146 (2012).

10. Watanabe, M., Uchida, K., Nakagaki, K., Trapnell, B. C. & Nakata, K. High avidity cytokine autoantibodies in health and disease: pathogenesis and mechanisms. Cytokine Growth Factor Rev. 21, 263–273 (2010).

11. Tabuchi, Y. et al. Protective effect of naturally occurring anti-HER2 autoantibodies on breast cancer. Breast Cancer Res. Treat. 157, 55–63 (2016).

12. Gillissen, M. A. et al. Patient-derived antibody recognizes a unique CD43 epitope expressed on all AML and has antileukemia activity in mice. Blood Adv 1, 1551–1564 (2017).

13. von Mensdorff-Pouilly, S. et al. Survival in early breast cancer patients is favorably influenced by a natural humoral immune response to polymorphic epithelial mucin. J. Clin. Oncol. 18, 574–583 (2000).

14. Naparstek, Y. & Plotz, P. H. The Role of Autoantibodies in Autoimmune Disease. (2003) doi:10.1146/annurev.iy.11.040193.000455.

15. Larman, H. B. et al. Autoantigen discovery with a synthetic human peptidome. Nat. Biotechnol. 29, 535–541 (2011).

16. Larman, H. B. et al. PhIP-Seq characterization of autoantibodies from patients with multiple sclerosis, type 1 diabetes and rheumatoid arthritis. J. Autoimmun. 43, 1–9 (2013).

17. Benjamin Larman, H., et al. Cytosolic 5′-nucleotidase 1A autoimmunity in sporadic inclusion body myositis: cN1A Autoimmunity in IBM. Ann. Neurol. 73, 408–418 (2013).

18. Meyer, S. et al. AIRE-Deficient Patients Harbor Unique High-Affinity Disease-Ameliorating Autoantibodies. Cell 166, 582–595 (2016).

19. Landegren, N. et al. Proteome-wide survey of the autoimmune target repertoire in autoimmune polyendocrine syndrome type 1. Sci. Rep. 6, 20104 (2016).

20. Fishman, D. et al. Autoantibody Repertoire in APECED Patients Targets Two Distinct Subgroups of Proteins. Front. Immunol. 8, 976 (2017).

21. Vazquez, S. E. et al. Identification of novel, clinically correlated autoantigens in the monogenic autoimmune syndrome APS1 by proteome-wide PhIP-Seq. Elife 9, (2020).

22. Kamath, K. et al. Antibody epitope repertoire analysis enables rapid antigen discovery and multiplex serology. Sci. Rep. 10, 5294 (2020).

23. Chen, W. S. et al. Autoantibody Landscape in Patients with Advanced Prostate Cancer. Clin. Cancer Res. 26, 6204–6214 (2020).

24. Laver, W. G., Air, G. M., Webster, R. G. & Smith-Gill, S. J. Epitopes on protein antigens: misconceptions and realities. Cell 61, 553–556 (1990).

25. Gai, S. A. & Wittrup, K. D. Yeast surface display for protein engineering and characterization. Curr. Opin. Struct. Biol. 17, 467–473 (2007).

26. Weiskopf, K. et al. Engineered SIRPα variants as immunotherapeutic adjuvants to anticancer antibodies. Science 341, 88–91 (2013).

27. Warren, J. T. et al. Manipulation of receptor oligomerization as a strategy to inhibit signaling by TNF superfamily members. Sci. Signal. 7, ra80 (2014).

28. Schweickhardt, R. L., Jiang, X., Garone, L. M. & Brondyk, W. H. Structure-Expression Relationship of Tumor Necrosis Factor Receptor Mutants That Increase Expression*. J. Biol. Chem. 278, 28961–28967 (2003).

29. Jin, M. et al. Directed evolution to probe protein allostery and integrin I domains of 200,000-fold higher affinity. Proc. Natl. Acad. Sci. U. S. A. 103, 5758–5763 (2006).

30. Chen, T. F., de Picciotto, S., Hackel, B. J. & Wittrup, K. D. Engineering fibronectin-based binding proteins by yeast surface display. Methods Enzymol. 523, 303–326 (2013).

31. Xu, G., Tasumi, S. & Pancer, Z. Yeast surface display of lamprey variable lymphocyte receptors. Methods Mol. Biol. 748, 21–33 (2011).

32. Chao, G., Cochran, J. R. & Wittrup, K. D. Fine epitope mapping of anti-epidermal growth factor receptor antibodies through random mutagenesis and yeast surface display. J. Mol. Biol. 342, 539–550 (2004).

33. Jeong, M.-Y., Rutter, J. & Chou, D. H.-C. Display of Single-Chain Insulin-like Peptides on a Yeast Surface. Biochemistry 58, 182–188 (2019).

34. Levin, A. M. et al. Exploiting a natural conformational switch to engineer an interleukin-2 “superkine.” Nature 484, 529–533 (2012).

35. Zhou, T. et al. IL-18BP is a secreted immune checkpoint and barrier to IL-18 immunotherapy. Nature 583, 609–614 (2020).

36. Ho, C. C. M. et al. Decoupling the Functional Pleiotropy of Stem Cell Factor by Tuning c-Kit Signaling. Cell 168, 1041–1052.e18 (2017).

37. Boder, E. T., Bill, J. R., Nields, A. W., Marrack, P. C. & Kappler, J. W. Yeast surface display of a noncovalent MHC class II heterodimer complexed with antigenic peptide. Biotechnol. Bioeng. 92, 485–491 (2005).

38. Birnbaum, M. E. et al. Deconstructing the peptide-MHC specificity of T cell recognition. Cell 157, 1073–1087 (2014).

39. Kieke, M. C. et al. Selection of functional T cell receptor mutants from a yeast surface-display library. Proc. Natl. Acad. Sci. U. S. A. 96, 5651–5656 (1999).

40. Rhiel, L. et al. REAL-Select: full-length antibody display and library screening by surface capture on yeast cells. PLoS One 9, e114887 (2014).

41. Boder, E. T. & Wittrup, K. D. Yeast surface display for screening combinatorial polypeptide libraries. Nat. Biotechnol. 15, 553–557 (1997).

42. Constantine, G. M. & Lionakis, M. S. Lessons from primary immunodeficiencies: Autoimmune regulator and autoimmune polyendocrinopathy-candidiasis-ectodermal dystrophy. Immunol. Rev. 287, 103–120 (2019).

43. Meager, A. et al. Anti-interferon autoantibodies in autoimmune polyendocrinopathy syndrome type 1. PLoS Med. 3, e289 (2006).

44. Wolff, A. S. B. et al. Autoimmune polyendocrine syndrome type 1 in Norway: phenotypic variation, autoantibodies, and novel mutations in the autoimmune regulator gene. J. Clin. Endocrinol. Metab. 92, 595–603 (2007).

45. Meloni, A. et al. Autoantibodies against type I interferons as an additional diagnostic criterion for autoimmune polyendocrine syndrome type I. J. Clin. Endocrinol. Metab. 93, 4389–4397 (2008).

46. Puel, A. et al. Autoantibodies against IL-17A, IL-17F, and IL-22 in patients with chronic mucocutaneous candidiasis and autoimmune polyendocrine syndrome type I. J. Exp. Med. 207, 291–297 (2010).

47. Kisand, K. et al. Chronic mucocutaneous candidiasis in APECED or thymoma patients correlates with autoimmunity to Th17-associated cytokines. J. Exp. Med. 207, 299–308 (2010).

48. Burbelo, P. D. et al. Profiling Autoantibodies against Salivary Proteins in Sicca Conditions. J. Dent. Res. 98, 772–778 (2019).

49. St-Pierre, C., Trofimov, A., Brochu, S., Lemieux, S. & Perreault, C. Differential Features of AIRE-Induced and AIRE-Independent Promiscuous Gene Expression in Thymic Epithelial Cells. J. Immunol. 195, 498–506 (2015).

50. Ferré, E. M. N. et al. Lymphocyte-driven regional immunopathology in pneumonitis caused by impaired central immune tolerance. Sci. Transl. Med. 11, (2019).

51. Lowe, M. E. STRUCTURE AND FUNCTION OF PANCREATIC LIPASE AND COLIPASE. (2003) doi:10.1146/annurev.nutr.17.1.141.

52. Tsokos, G. C., Lo, M. S., Costa Reis, P. & Sullivan, K. E. New insights into the immunopathogenesis of systemic lupus erythematosus. Nat. Rev. Rheumatol. 12, 716–730 (2016).

53. Pisetsky, D. S. & Lipsky, P. E. New insights into the role of antinuclear antibodies in systemic lupus erythematosus. Nat. Rev. Rheumatol. 16, 565–579 (2020).

54. Bombardier, C., Gladman, D. D., Urowitz, M. B., Caron, D. & Chang, C. H. Derivation of the SLEDAI. A disease activity index for lupus patients. The Committee on Prognosis Studies in SLE. Arthritis Rheum. 35, 630–640 (1992).

55. Yang, Z., Liang, Y., Xi, W., Li, C. & Zhong, R. Association of increased serum IL-33 levels with clinical and laboratory characteristics of systemic lupus erythematosus in Chinese population. Clin. Exp. Med. 11, 75–80 (2011).

56. Guo, J. et al. The association of novel IL-33 polymorphisms with sIL-33 and risk of systemic lupus erythematosus. Mol. Immunol. 77, 1–7 (2016).

57. Rose, W. A., 2nd et al. Interleukin-33 Contributes Toward Loss of Tolerance by Promoting B-Cell-Activating Factor of the Tumor-Necrosis-Factor Family (BAFF)- Dependent Autoantibody Production. Front. Immunol. 9, 2871 (2018).

58. Li, P., Lin, W. & Zheng, X. IL-33 neutralization suppresses lupus disease in lupus-prone mice. Inflammation 37, 824–832 (2014).

59. Herscovics, A. & Orlean, P. Glycoprotein biosynthesis in yeast. FASEB J. 7, 540–550 (1993).

60. Hamilton, S. R. & Gerngross, T. U. Glycosylation engineering in yeast: the advent of fully humanized yeast. Curr. Opin. Biotechnol. 18, 387–392 (2007).

61. Picelli, S. et al. Tn5 transposase and tagmentation procedures for massively scaled sequencing projects. Genome Res. 24, 2033–2040 (2014).

62. Robinson, M. D., McCarthy, D. J. & Smyth, G. K. edgeR: a Bioconductor package for differential expression analysis of digital gene expression data. Bioinformatics 26, 139–140 (2010).

63. Petri, M. et al. Derivation and validation of the Systemic Lupus International Collaborating Clinics classification criteria for systemic lupus erythematosus. Arthritis Rheum. 64, 2677–2686 (2012).

64. Ferre, E. M. N. et al. Redefined clinical features and diagnostic criteria in autoimmune polyendocrinopathy-candidiasis-ectodermal dystrophy. JCI Insight 1, (2016).

